# Default mode network spatio-temporal electrophysiological signature and causal role in creativity

**DOI:** 10.1101/2023.09.13.557639

**Authors:** E. Bartoli, E. Devara, H.Q. Dang, R. Rabinovich, R.K. Mathura, A. Anand, B.R. Pascuzzi, J. Adkinson, K.R. Bijanki, S.A. Sheth, B. Shofty

## Abstract

The default mode network (DMN) is a widely distributed, intrinsic brain network thought to play a crucial role in internally-directed cognition. It subserves self-referential thinking, recollection of the past, mind wandering, and creativity. Knowledge about the electrophysiology underlying DMN activity is scarce, due to the difficulty to simultaneously record from multiple distant cortical areas with commonly-used techniques. The present study employs stereo-electroencephalography depth electrodes in 13 human patients undergoing monitoring for epilepsy, obtaining high spatiotemporal resolution neural recordings across multiple canonical DMN regions. Our results offer a rare insight into the temporal evolution and spatial origin of theta (4-8Hz) and gamma signals (30-70Hz) during two DMN-associated higher cognitive functions: mind-wandering and alternate uses. During the performance of these tasks, DMN activity is defined by a specific pattern of decreased theta coupled with increased gamma power. Critically, creativity and mind wandering engage the DMN with different dynamics: creativity recruits the DMN strongly during the covert search of ideas, while mind wandering displays the strongest modulation of DMN during the later recall of the train of thoughts. Theta band power modulations, predominantly occurring during mind wandering, do not show a predominant spatial origin within the DMN. In contrast, gamma power effects were similar for mind wandering and creativity and more strongly associated to lateral temporal nodes. Interfering with DMN activity through direct cortical stimulation within several DMN nodes caused a decrease in creativity, specifically reducing the originality of the alternate uses, without affecting creative fluency or mind wandering. These results suggest that DMN activity is flexibly modulated as a function of specific cognitive processes and supports its causal role in creative thinking. Our findings shed light on the neural constructs supporting creative cognition and provide causal evidence for the role of DMN in the generation of original connections among concepts.

## Introduction

Internal mentation and other introspective thought processes underlie the stream of human consciousness. The default mode network (DMN), a fundamental neurobiological system, is a dispersed cortical network that is thought to underlie spontaneous cognitive processes (Buckner et al 2008). While prominent during periods of rumination or even at rest, the DMN deactivates when attention is directed to the external world and towards other cognitively demanding tasks (Raichle et al 2001, Shulman et al 1997). The discovery of DMN using fMRI has led to a plethora of functional imaging-based studies correlating DMN activity with various cognitive functions and disease states (Buckner 2012, Buckner & DiNicola 2019, Mohan et al 2016). Some of these studies have emphasized activity within the DMN, investigating activation during cognitive processes such as episodic memory retrieval, future simulation, and mind wandering, while others have looked at DMN deactivation in response to externally directed tasks in the form of visual search or mental arithmetic (Addis et al 2007, Christoff et al 2009, Fox et al 2015, Svoboda et al 2006).

Despite the proliferation of neuroimaging studies that have queried the human DMN over the past twenty years, there still exists a relative paucity of information regarding the electrophysiological underpinnings of the network (Buckner & DiNicola 2019). The use of intracranial electroencephalography (iEEG) to study the DMN and other wide-spread cortical networks have gained momentum over the last few years. The high temporal resolution obtained through cortical recordings, as well as the precise anatomical localization of the electrodes, allows for in-depth investigation into network-based function and the underlying, millisecond neurophysiology (Das et al 2022, Fox et al 2018). The clinical transition from a surface-based investigation of intractable focal epilepsy (via cortical grids and strips) to minimally invasive, spatially dispersed sampling technique (via depth electrodes for stereo-EEG, sEEG) further extends our ability to decipher the neural mechanisms underlying higher cognitive functions. Indeed, studies employing surface cortical grids and strips were unlikely to record from more than one of the several areas comprising the DMN due to the network’s dispersed spatial structure, while sEEG trajectories now offer access to a wider array of distant regions, including mesial structures such as the cingulate and medial prefrontal cortices, within the same subject. For this reason, most of the pioneering studies on the electrophysiological basis of the DMN have been focused on individual hubs. Earlier studies looking into intracranial electrophysiology of the DMN addressed mechanistic network properties and their correlation with previous fMRI BOLD findings, providing essential evidence matching fMRI activation and deactivation with massive neuronal firing patterns as identified using local field potentials (Keller et al 2013). Most of these studies have centered on investigations into high-frequency neural signals, known to be a robust correlate of fMRI blood oxygen level-dependent (BOLD) signals and, to a given extent, spiking activity in the brain (Leszczynski et al 2020, Ray et al 2008).

Localized deactivations in high-frequency activity in default mode regions such as the posterior cingulate cortex (PCC), ventrolateral and rostromedial prefrontal cortex have been noted on a variety of externally focused tasks, including mental arithmetic, visual search, and reading (Foster et al 2012, Foster et al 2015, Jerbi et al 2010, Jung et al 2010, Lachaux et al 2008, Ossandon et al 2011). Conversely, task-induced elevations in high-frequency activity within the DMN have been discussed in the context of internally directed cognition, reported in areas such as the PCC and retrosplenial cortex in tasks of self-referential judgement and autobiographical memory (Daitch & Parvizi 2018, Dastjerdi et al 2011, Foster et al 2013, Foster & Parvizi 2012). More recent investigations started to unravel the correspondence of inter- and intra-network dynamics with cognitive functions, leveraging the aforementioned advantages of sEEG recordings to dissect the network’s role in complex behavior. For example, mentalizing about self and others recruits the DMN with a specific posterior to anterior spatio-temporal pattern (Tan et al 2022). Within DMN areas, correlated slow fluctuations (<4Hz) occur during rest, memory encoding and recall, while cross-network interactions seem to rely on higher frequency-band signals (Das et al 2022). These types of intra-network slow fluctuations at rest have been shown to modify firing rates and gamma-band (>30Hz) local field potentials (Nir et al 2008) and are a reliable correlate of BOLD-signal spontaneous fluctuations (Kucyi et al 2018). Inter-network interactions have been reported in the beta (13-20Hz) and in the theta range (4-8Hz) (Das et al 2022, Foster et al 2013) during memory encoding/recall and autobiographical memory retrieval respectively. Despite the substantial amount of evidence indicating the involvement and interaction of the DMN with other networks during various complex cognitive functions, there is very limited causal evidence. A large-sample study employing direct stimulation to interfere with posteromedial cortex activity at rest, did not produce any notable effects (Foster & Parvizi 2017). However, a recent study demonstrated a causal link between creative thinking and DMN integrity using individualized direct cortical stimulation during task performance (Shofty et al 2022). Disrupting the network caused a reduction in the number of creative responses during an alternate-uses task (i.e., requiring to formulate non-canonical uses for everyday objects). Indeed, DMN may play a critical role in creative cognition by supporting divergent thinking, in line with its distinctive role in internal mental manipulation and simulation (Jung et al 2013, Mok 2014). From this perspective, creative thinking is a high-order DMN-associated function that requires retrieving and combining several different types of information to generate a novel and useful idea.

Though significant strides have been made, we have yet to assess the full span of spatiotemporal dynamics occurring within the DMN as a function of the ongoing cognitive processes. How does the DMN reorganize and modify its activity based on different modes of thought? What are the neural signals underlying these dynamics? What parts of the network are flexibly recruited to support its various cognitive domains? Is the DMN necessary for creative thinking? In the present study, we examine two different DMN-associated thought processes, mind wandering and creativity, in a sample of patients undergoing monitoring with intracranial electrodes as part of their epilepsy evaluation. Taking advantage of the distributed spatial coverage and high temporal resolution of sEEG recordings, we were able to precisely record neural activity within the canonical DMN during these cognitive tasks. In addition, we leveraged electrical high-frequency stimulation to disrupt DMN integrity and test the causal role of the DMN in both mind wandering and creativity. Through these experiments, we link unique patterns of theta and gamma range signals to different stages of mind wandering and creativity. In addition, we demonstrate that disrupting DMN activity reduces creativity without affecting mind wandering.

## Materials and Methods

### Human subjects

Thirteen subjects (7 males, 6 females, mean age of 41 years, ranging from 19-60 years) consented to participate in this study while they underwent invasive epilepsy monitoring with stereo-electroencephalography (sEEG) at Baylor St. Luke’s Medical Center (Houston, Texas, USA). Detailed subject information is reported in Table 1. Experimental procedures approved by the Institution Review Board at Baylor College of Medicine (IRB protocol number H-18112). No patients with anatomical abnormalities, or prior surgical resection in the areas of interest participated in this study. Experiments were recorded while inter-ictal epileptic discharges were absent in the areas of interest. Stimulation was performed when deemed safe by the clinical team supervising the experiment.

**Table 1.** Sample information. For each participant in the study (n=13) the table reports the age at the time of the experiment (years), the sex (M=male, F=female), the number of electrodes in dorso-medial and lateral temporal-DMN and their sum (count), whether the subject took part in the stimulation experiment (yes/no), and the stimulation location (R L ACC: right or left anterior cingulate; MTG: middle temporal gyrus; SFG-dLPFC superior frontal gyrus extending into dorso-lateral prefrontal cortex)

| # Subject | Age | Sex | Electrodes in dorsomedial-DMN | Electrodes in lateral temporal-DMN | Total Electrodes in DMN | Stimulation Experiment | Stimulated locations |
| --- | --- | --- | --- | --- | --- | --- | --- |
| 1 | 47 | M | 14 | 13 | 27 | y | RACC |
| 2 | 19 | M | 8 | 27 | 35 | n |  |
| 3 | 57 | M | 8 | 8 | 16 | y | LACC |
| 4 | 24 | M | 16 | 21 | 37 | y | LACC |
| 5 | 52 | F | 16 | - | 16 | y | RACC |
| 6 | 53 | F | - | 9 | 9 | n |  |
| 7 | 25 | F | 11 | 11 | 22 | y | MTG |
| 8 | 41 | F | 6 | 24 | 30 | y | SFG-dLPFC |
| 9 | 60 | M | 6 | - | 6 | n |  |
| 10 | 44 | F | 3 | 20 | 23 | y | LACC |
| 11 | 46 | F | 5 | 23 | 28 | n |  |
| 12 | 42 | M | 5 | 19 | 24 | y | SFG-dLPFC |
| 13 | 25 | M | 50 | 10 | 60 | y | RACC |

### Task design

All subjects performed the experimental paradigm while reclined in a hospital bed in a quiet room. All tasks were presented on an adjustable monitor (1920×1080 resolution, 47.5×26.7 cm screen size, 60Hz refresh rate, connected to a PC running Windows 10 Pro at a viewing distance of 57 cm such that 1 cm = ∼1 degree visual angle). Tasks were programed using Psychtoolbox-3 functions (v3.0.16) (Brainard 1997) running on MATLAB (R2019a, MathWorks, MA, USA). An experimenter was seated by the bedside throughout the experiment to give subjects instructions. The experiment consisted of three types of tasks (Fig. 1): a mind wandering task (MW), an alternate uses task (AUT), and a visual attention task (ATT). During MW, the subjects fixated on a colored shape in the center of the screen for 20 seconds. They were then asked, “What were you thinking about just now?” to which they verbalized their thoughts for as long as they could or until 1 minute had passed. The MW task was adapted from the “Shape Expectations” task developed by O’Callaghan and colleagues (O’Callaghan et al 2015). During AUT, the subjects fixated on an object in the center of the screen for 20 seconds and were informed of its typical function. They were then asked, “What are other uses for this object?” to which they listed out as many alternate uses that they could conceive or until 1 minute had passed. Their responses to the MW and AUT tasks were recorded verbatim into Microsoft Excel (Microsoft, WA, USA). During ATT, the subjects attended to the center of the screen where a fixation cross would flash (250 ms) to signal the start of the ATT trial: subjects were required to attend the screen for 400 ms and report the detection of a target stimulus (small white disk, subtending 0.5 degrees visual angle) by button press (max response time allowed 2 seconds). The target would briefly flash on 75% of trials (for a single frame, ∼17 ms), either on its own or followed by a white annulus mask (1 degree, flashed for 2 frames, with a 50ms lag between target and mask). On the remaining 25% trials only the annulus mask appeared without the target. Button presses were recorded by the experiment software. Between each trial there was a short break (gray screen, 2 seconds). The experiment was divided into three blocks each comprised of 4 MW trials, 4 AUT trials and 8 ATT trials (16 total trials per block). For the MW and AUT trials, different colored shapes and objects, respectively, were used across all blocks such that no stimuli were repeated to avoid learning effects. The stimuli used in each task were identical for all patients and their order was randomized within each block. A subset of patients (n=9) received intracranial stimulation during a portion of the experiment (block 2).

**Figure 1.**
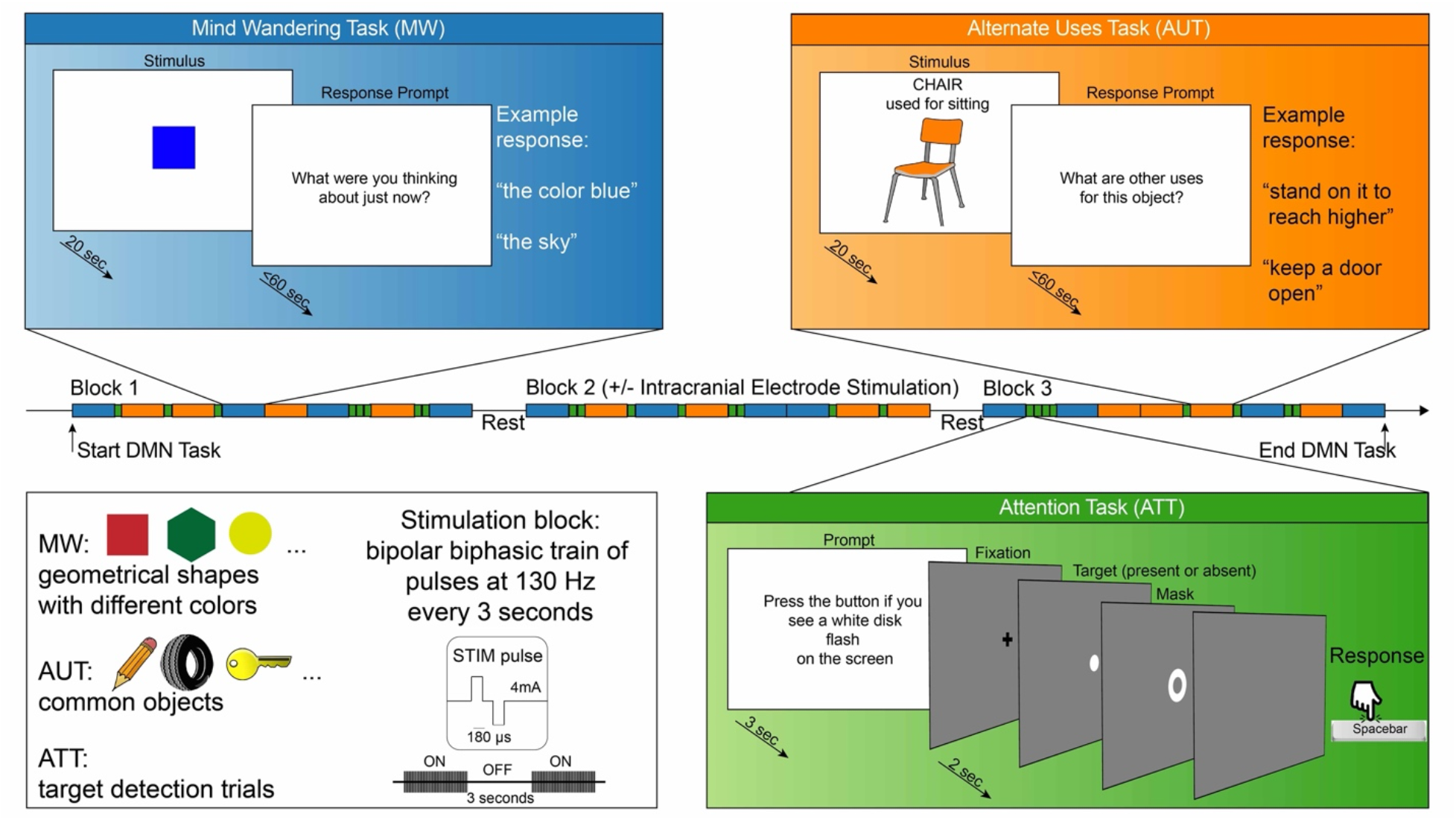
Experimental Task Design. Each box shows a schematic of one of the three tasks: mind wandering, alternate uses, and attention (MW, AUT, ATT) plus an overview of the stimuli and the stimulation parameters. In the middle, a schematic depicts the subdivision of the experiments in three blocks, with stimulation being delivered during the second block. The MW and AUT trials had a stimulus presentation stage and a response stage. During the stimulus stage a visual cue indicated to let the mind freely wander (MW) or to think about alternative uses for the displayed object (AUT). During the response stage, the participant was probed to verbalize either the train of thoughts that just occurred (MW) or to list the alternative uses for the item just displayed (AUT). In the attention task, the subject was cued to detect the brief presentation of a white disk, which could appear alone (target only), followed by a white ring mask (target plus mask), or not appear at all (mask only).

### sEEG probes

Reported data were acquired by sEEG depth probes. The sEEG probes had either a 0.8 mm diameter, with 8 to 16 electrode contacts along the probe with a 3.5 mm center-to-center distance (PMT Corporation, MN, USA) or a 1.28 mm diameter, with 9 recording contacts, with a 5.0 mm center-to-center distance between contacts (AdTech Medical Instrument Corporation, WI, USA).

### Electrode localization and selection

For each subject, we determined electrode locations by employing the software pipeline intracranial Electrode Visualization, iELVis (Groppe et al 2017). In short, the post-operative CT image was registered to the pre-operative T1 anatomical MRI image using FSL (Jenkinson et al 2012). Next, the location of each electrode was identified in the CT-MRI overlay using BioImage Suite (Papademetris et al 2006). The electrode anatomical locations were classified based on their proximity to the cortical surface model, reconstructed by the T1 image using Freesurfer (version 6.0; (Dale et al 1999)). The anatomical assignment of each electrode was verified by an expert in neuroanatomy (B.S.). We further labelled each electrode by mapping onto each individual cortical surface the 7 Network estimate atlas (Yeo et al., 2011) and finding the most likely cortical parcellation estimate using a 5 mm radius around each electrode. The atlas consists of 7 networks based on intrinsic connectivity. To visualize electrodes of interest on a common brain, we transformed electrode coordinates into the MNI average stereotaxic model and represented them as spheres on the Freesurfer Colin27 brain using Multi-Modality Visualization Tool, MMVT (Felsenstein et al 2019) (Figure 2A). Electrodes classified as recording from the Default mode network (DMN) were selected for the current study. We further identified two subsystems within the DMN: medial and dorso-lateral frontal-parietal areas, which we labeled dorsomedial-DMN (electrodes recording from ventromedial prefrontal cortex, anterior cingulate, superior frontal gyrus, posterior cingulate, posterior parietal, parahippocampal gyrus), and lateral temporal areas, which we labeled lateral temporal-DMN (electrodes recording from middle temporal gyrus, superior and middle temporal sulci). Note that the division into these two subsystems is partially dictated by the cortical sampling availability. In particular, the dorsomedial-DMN system includes both frontal, cingulate and parietal contributions and ideally it would have been interesting to isolate each unique contribution, but our sample did not provide enough within-subject cortical coverage across these areas to reliably interpret any subdivision beyond the one presented here.

**Figure 2.**
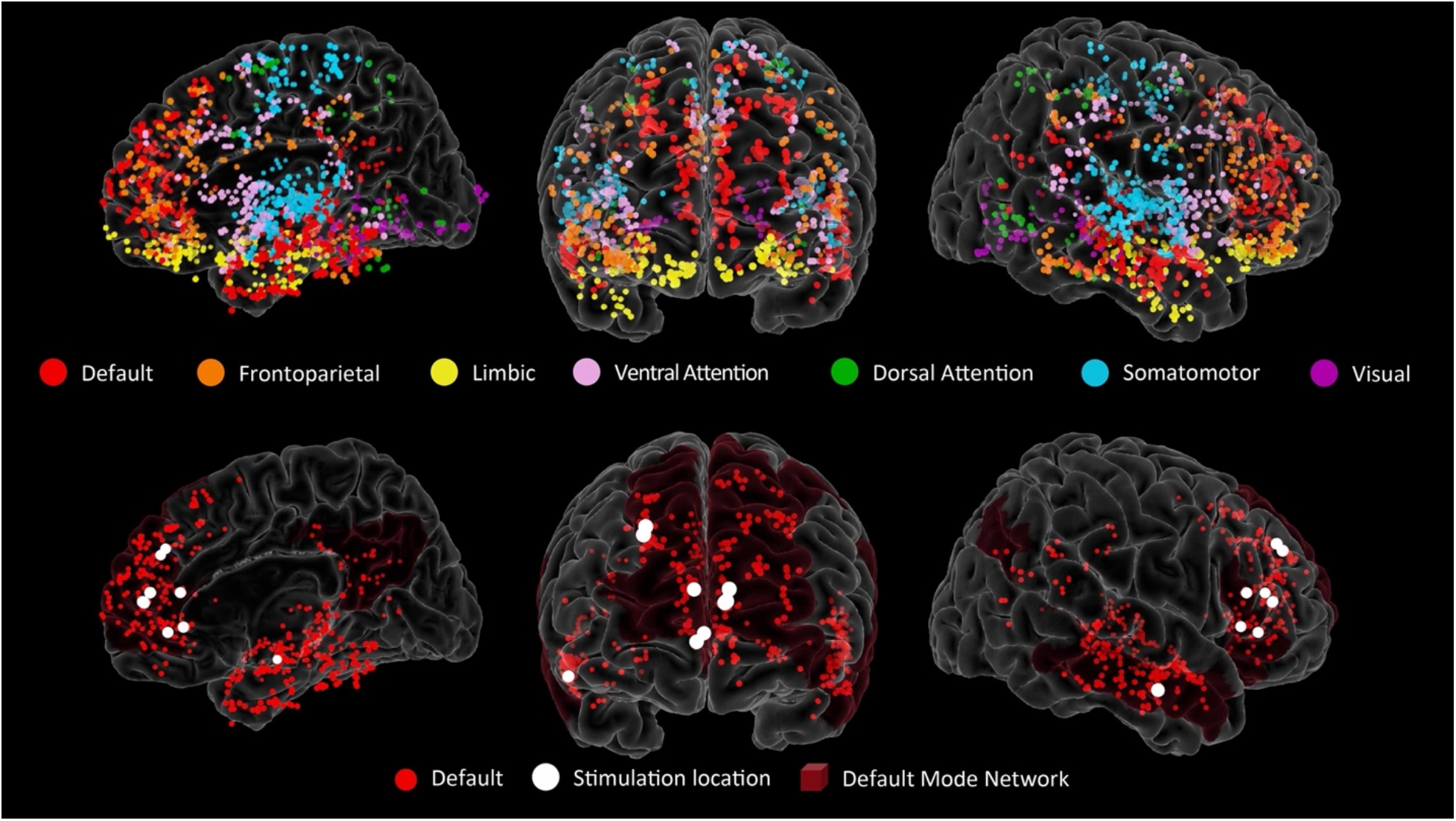
Electrode locations. Top row shows all the intracranial electrode locations (13 subjects) represented as spheres on a template brain, color coded by the Yeo Atlas assignment based on their proximity to the cortical surface. Bottom row shows only the electrodes the DMN network (red) and the location of the stimulation contacts (white). The stimulation electrodes are represented as a larger sphere centered between the pair of electrodes used for bipolar stimulation, 9 subjects: 1 right middle temporal gyrus, 2 right frontal superior gyrus, 3 right anterior cingulate, 3 left anterior cingulate; note that 2 of the 3 locations in the left anterior cingulate are almost completely overlapping).

### Electrophysiological recording and preprocessing

sEEG signals were recorded by a 256 channel BlackRock Cerebus system (BlackRock Microsystems, UT, USA) at 2kHz sampling rate, with a 4^th^ order Butterworth bandpass filter of 0.3-500Hz. sEEG recordings were referenced to an electrode contact visually determined to be in white matter. A photoresistor sensor recording at 30kHz was attached to the task monitor to synchronize intracranial recordings to the stimulus presentation time. All signals were processed by custom scripts in MATLAB (R2019a, MathWorks, MA, USA). Raw sEEG signals were firstly inspected for line noise, recording artifacts, and interictal epileptic spikes. Electrodes with artifacts and epileptic spikes were excluded from further analysis. After this exclusion was applied, we obtained 333 total electrodes in the DMN (148 electrodes in dorsomedial-DMN and 185 in lateral-DMN), ranging from 6-60 for each subject. Across our sample, 10 subjects had electrodes recording from both DMN subsystems, 2 subjects only from dorsomedial-DMN and 1 subject only from lateral-DMN. Next, the signal from each electrode was notch filtered (60 Hz and harmonics) and re-referenced to the neighboring electrodes using a Laplacian local average reference. Finally, re-referenced signals were downsampled to 500 Hz and spectrally decomposed using a family of Morlet wavelets, with center frequencies ranging logarithmically from 1 to 200 Hz in 100 steps (7 cycles).

### Electrophysiological stimulation

Bipolar stimulation (trains of pulses at 130 Hz, with 4 mA amplitude and 180 us pulse width) was delivered in cycles of 3 seconds on, 3 seconds off to a pair of electrode contacts located within one DMN node, using a Cerestim R96 stimulator (BlackRock Microsystems, UT, USA). Electrophysiological signals were not analyzed during stimulation. Stimulation occurred during the entire second block of the experiment. Out the 13 participants, 4 participants did not receive stimulation due to clinical factors (i.e. timing considerations, seizure onset zone proximity, etc.). Two patients received stimulation with modified parameters (Subject#1: 50Hz; Subject#4: 90 us pulse width). For each of the 9 participants receiving stimulation, a pair of electrode contacts within the DMN network was selected (3 in left ACC, 3 in right ACC, 1 in right MTG, 2 in right SFG-dlPFC; Fig. 2B). The electrodes were selected based on anatomical and clinical considerations (not part of the suspected seizure onset zone) and were subjected to an established pipeline to evaluate the safety of the stimulation targets and parameters with respect to epileptiform activity (Goldstein et al 2019). Stimulation was delivered throughout block 2 and the electrophysiological data from the stimulation period was excluded from the analysis.

### Behavioral Scoring

The AUT and MW tasks were scored based on the semantic distance between each object or item (e.g. chair; blue square) and the responses that the subjects conceived. Semantic distance captures the relationship between ideas, concepts, or other texts, based on the assumption that words that occur in similar contexts are similar in meaning (Gunther et al 2019). The response data were collated into a Microsoft Excel sheet individually for each patient. Each instance of the original object was listed next to a single corresponding response. Responses that did not make sense in the context of the task were removed from the dataset. Scoring the originality of the responses was performed using SemDis, an automated algorithm to compute semantic distance developed by Beaty and Johnson, and available at http://semdis.wlu.psu.edu/ (Beaty & Johnson 2021). SemDis scores range from 0-2 with increasing score indicating greater semantic distance. This algorithm not only combines multiple commonly used methods to vectorize and calculate the cosine distance between resulting vectors, but also considers the meaning of the words in multiple semantic spaces. This process results in a more accurate representation of the relationship between the words/phrases than any individual method. Each patient’s behavioral data was uploaded into the SemDis website, using the following steps: 1) remove filler words (e.g., a, the, and), leaving only the words that represent the true meaning of the phrase; 2) clean the data, removing numbers, symbols, and other special characters that cannot be vectorized; 3) use a multiplicative compositional model; 4) calculate the SemDis scores in all five available semantic spaces to fully capture various meanings of both the cues and responses; and 5) use the average score across all semantic spaces to compute the mean SemDis score for each item.

For further behavioral analysis (to score variability and flexibility of responses), we used the spaCy python library (Honnibal & Montani 2017) to perform the following steps. We first cleaned the phrases by removing punctuation and stop words (spaCy defaults, as well as the words “use”, “things”, and “stuff”). Then, we vectorized the phrases using the Universal Sentence Encoder text embedding model (Cer et al 2018). To calculate variability between a set of responses for a given item, we determined the semantic distance between each pair of responses by calculating the cosine distance between the vectors corresponding to those phrases; we then used the sum of these distances (i.e., the total cosine distance) as a measure of variability. To investigate how responses clustered into discrete categories, we used Python’s scikit-learn software (Pedregosa et al 2011) to perform t-SNE clustering (parameters: perplexity = 5; learning rate = 1) on all patients’ responses for each item. As a result of this dimensionality reduction process, we were able to visualize the responses in 2 dimensions. Subsequently, we used the scikit-learn affinity propagation clustering algorithm to identify semantic categories.

Finally, the ATT task served to track whether the subjects were engaged and paying attention to the screen. Button presses (corresponding to participants reporting a target detection) were tabulated by the experiment software and stored in a behavioral file. The probability of reporting a target (hit rate) was higher for target only trials than for trials with the target followed by the mask (70% and 43%), the false alarm rate (reporting a target during mask only trials) was 20%. This pattern confirmed that brief duration of the target and the backward masking made the attentional task challenging, and the 70% hit rate to target only trials confirmed that the participants were engaged and paying attention. The task also served as a control for the specificity of the results for default versus attentional cognitive domains.

### Statistical analyses

Statistical analyses focused on changes in signals in the theta and gamma frequency ranges. Frequency band power was obtained by averaging the magnitude of the Morlet wavelet decomposition result between 4-8 Hz for theta and 30-70 Hz for gamma. Power values during each trial were normalized to percent change with respect to the pre-trial baseline period (500-100 ms before stimulus onset). Power % change values computed for each electrode location were then averaged over trials. Primarily, multi-level mixed effects models were adopted to account for the unbalanced and nested data structure in the current study (i.e., multiple electrodes from each subject): experimental conditions were set as fixed effects, while subjects and electrodes were set as nested random effects. This approach accounts for the non-independence between observations (e.g., different electrodes from the same subject) while preserving the richness of the dataset (e.g., avoiding averaging across electrodes). In practice, to evaluate the importance of a fixed effect (e.g. task type) the model was compared to a simpler model without the coefficient of interest but with identical random effect structure using Chi-square difference tests. This approach tests if the more complex model is significantly better at capturing the data (i.e. if the added parameters yield to a significant reduction of the residuals). Post-hoc differences were evaluated by using z-statistic approximation with two-sided probability and adjusting p-values for multiple comparison using Bonferroni (indicated by p-adj in the results). Exact p-values are reported for values larger than 0.001 (any smaller p-values are simplified as p<0.001). All analysis were performed in R (R Development Core Team 2010) using lmer (Bates et al 2014) and multcomp packages.

#### DMN engagement

First, we compared the neural signatures of the DMN occurring during the viewing of the stimuli for the three different tasks: AUT, MW, and ATT. By nature of the task, each trial of ATT takes place over a much shorter time window than AUT or MW. To avoid confounds related to the different temporal duration of attention and default mode tasks (AUT and MW), we focused on the initial engagement of the DMN in a common time window (2 seconds), fully capturing the attentional task (ATT) and the initial encoding of the object (AUT) or shape (MW). We employed theta and gamma power values computed over the first 2 seconds from stimulus onset (employing a sliding 1 s window with 50% overlap). A linear mixed effects model for each frequency range of interest (i.e., theta and gamma) was utilized to assess variations in band power as a function of the task type (ATT, AUT, MW: fixed effect; subject and electrodes: nested random effects). We performed two control analysis: first, we repeated the analysis changing the time-window employed. We marked the shortest ATT trial in each patient and averaged theta and gamma power over the window between task onset and this identified time. This ensured that the data being analyzed for the attention task did not include button presses. Second, we repeated the analysis using electrodes not part of the DMN, selecting electrodes localized in the somatomotor network.

#### DMN dynamics during creativity and mind wandering

Next, we probed further into the DMN dynamics during the two default mode tasks, AUT and MW. As detailed previously, the tasks were comprised of two stages: a stimulus presentation stage, termed “Stimulus”, and a patient response stage, termed “Response”. We computed theta and gamma power during the first 15 seconds from the onset of each stage (employing a sliding 1 s window with 50% overlap, leading to 29 time-bins centered 500ms apart, starting at 500ms from the onset of each stage; e.g. the first time bin is centered at 500ms from the onset and computed by averaging the data from 0-1s, the second is centered at 1s computed over 0.5-1.5, etc.). The 15 second window was chosen as a trade-off to give ample time to capture creativity and mind-wandering activity in AUT and MW during both stages while capturing the relevant events. This was important as during the response stage the time of active response production/verbalization varied, lasting on average 32 seconds (25% percentile: 18 seconds, 75%: 45 seconds). Based on the distribution of the verbal response durations we selected 15 seconds as a compromise to capture meaningful events during both stages across trials and patients. Variations in band power were investigated with linear mixed effects models for theta and gamma, designating task type (AUT, MW), task stage (Stimulus, Response), time bin (0 to 15s in 500 ms steps) and DMN-subsystems (dorsomedial-DMN, lateral-DMN) as fixed effects, subjects and electrodes as nested random effects. The importance of each effect and interaction coefficient was assessed using sequential model comparisons (starting from a model with only task type as a fixed effect and adding the effect of task type, the interaction between the two effects, etc.) as described above.

#### Interplay between neural signatures and DMN nodes

To evaluate the interaction between theta and gamma power dynamics across the dorsomedial and lateral-DMN nodes and their similarity during the two tasks, we employed hierarchical clustering. This analysis was restricted to a subset of patients with electrodes in both DMN nodes (n=10). The distance between the variables task type, and DMN-node were computed using 1-correlation values between the observations formed by the theta and gamma power dynamics (0-15s in 500 ms steps) for both task stages (Stimulus and Response) concatenated for each subject (averaging across electrodes recording from the same DMN node within subject), leading to 580 observations for each variable (29 time points by 2 task stages by 10 subjects). We included the overall theta and gamma power (averaged across DMN nodes and tasks) as variables to identify the major contributors to the overall DMN engagement. The clustering was performed using complete-linkage and the reliability of the clusters statistically tested with bootstrap resampling (n=10,000) using pvclust and corrplot packages. Bootstrap probability (probability to obtain the same cluster over the resampling iterations) is reported to highlight the robustness of the result.

#### Stimulation effect on behavior

The MW and AUT creativity scores for all subjects taking part in the stimulation portion of the experiment (n=9) were computed for all stimuli (colored shape or object) and analyzed according to stimulation status (no stimulation vs. stimulation). The SemDis scores are bound between 0-2, with higher values representing higher semantic distance between the stimulus and the response, a proxy of originality. Given the non-normality of the SemDis score we employed non-parametric testing (Wilcoxon Signed Rank Test, using subject as a pairing variable) to test for the effect of stimulation status on the trial-averaged SemDis scores for both AUT and MW (i.e., averaging the sematic distance scores within stimulation status for each patient). For any metric or task demonstrating an effect of stimulation, we employ a control analysis on the full sample (n=13, i.e. including the 4 subjects that did not receive stimulation due to experimental constraints) and all trials (not averaging the scores across trials): non-parametric testing for differences between trials with stimulation versus trials without stimulation (Mann-Whitney Test, independent samples). This control analysis is used to evaluate if the difference between stimulation and no stimulation trials can be also detected when considering all subjects (and not only as a within-subject effect). The same approach was employed for other behavioral score metrics (variability, flexibility).

## Results

### DMN engagement at stimulus onset displays opposite theta and gamma modulations

Power in the theta and gamma frequency-bands in the DMN were differentially modulated during the viewing of the stimuli for the three different tasks (alternate uses task, AUT; mind wandering, MW; and attention task, ATT; Figure 3A). Both DMN theta and gamma were significantly modulated by the task type (theta model comparison without/with task type as a fixed effect: χ^2^(2)=89.04, p<0.001; gamma: χ^2^(2)=38.45, p<0.001). As a whole, low and high frequency power both increased during all three tasks. However, the alternate uses and mind wandering tasks were characterized by a *relative* decrease (i.e., smaller increase) in theta power compared to the attention task (mean ± standard error computed across subjects and electrodes; ATT: 173.8±3.9%; AUT: 123.6±4.01%; MW: 140.7±3.9%). Pairwise post-hoc comparisons confirmed that theta power differed between all three tasks, with the alternate uses task showing the strongest relative decrease (smallest increase) in DMN theta (AUT vs. ATT: z=-9.6, p-adj<0.001; MW vs. ATT: z=-6.3, p-adj<0.001; MW vs. AUT: z=3.3, p-adj=0.003).

**Figure 3.**
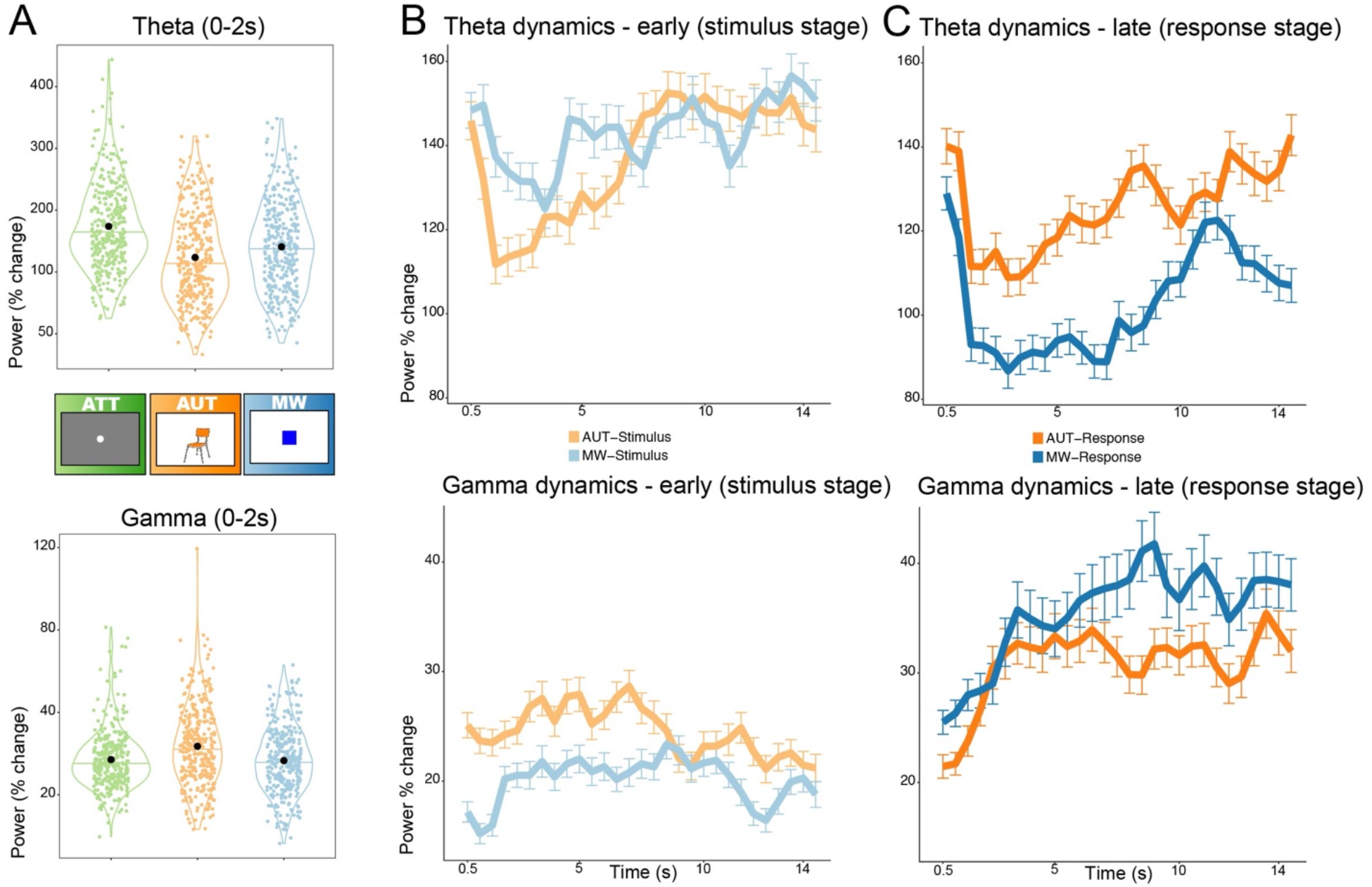
Engagement of DMN activity during the tasks. Panel A: Theta (4-8Hz, top) and gamma (30-70Hz, bottom) power modulations recorded in DMN during the first 2 seconds from stimulus onset for each task, as percent change from pre-stimulus baseline. All tasks increased theta and gamma power above baseline. Comparing the tasks, theta power was highest during attention (ATT) followed by mind wandering (MW) and was lowest for creativity (AUT), all comparisons p-adj<0.01. Gamma power was modulated in the opposite way, being largest for AUT and with no differences between MW and ATT. Each point in the violin plots represents an electrode in the DMN (n=333). Panel B: temporal evolution of theta and gamma activity during creativity (AUT, orange) and mind wandering (MW, blue) from stimulus onset (lighter colors) and Panel C: from response prompt (darker colors). The dynamics reveal how the DMN is engaged differently according to the task stage: during the first 6-8 seconds of viewing the stimuli, theta and gamma are more strongly modulated by AUT than MW. In the response stage, the opposite pattern emerges, with a stronger theta decrease and gamma increase occurring during the verbalization of the train of thoughts in MW.

In the gamma range, the opposite pattern emerged, with the alternate uses task inducing the largest increase in gamma power; mind wandering and attention both yielded lower increases in gamma power, and there was no significant difference between these two tasks (ATT: 17.1±0.79%; AUT: 23.5± 1.04%; MW: 16.6±0.87%; AUT vs. ATT: z=5.2, p-adj<0.001; MW vs. ATT: z=-0.4, p-adj=1; MW vs. AUT: z=-5.6, p-adj<0.001). The same analysis was repeated using a shorter time-window (based on the ATT trial length), yielding virtually identical results. Overall, by contrasting the initial stage of default mode tasks versus attention, we report that DMN engagement features a relative decrease in theta power for both AUT and MW, coupled with an increase in gamma-range activity occurring specifically for the alternate uses task (Figure 3A). As a control, we repeated the above analysis on data collected from electrodes in a different network (pre-central gyrus, somatomotor network). The theta activity pattern in the somatomotor network showed a task-related modulation different from the one found in the DMN. Specifically, theta power only increased for the attention task, and no differences were present between the default mode tasks (model comparison with/without fixed effect of task: χ^2^(2)=15.8, p<0.001; AUT vs. ATT: z=-4.2, p-adj<0.001; MW vs. ATT: z=-2.2, p-adj=0.08; MW vs. AUT: z=1.99, p-adj=0.14). Gamma activity in the somatomotor network was not modulated by any of the tasks, and no differences between tasks were found (adding the fixed effect of task type was not significantly improving the model, χ^2^(2)=0.38, p=0.8; all post-hoc comparisons between tasks not significant). Thus, the aforementioned differences in the theta and gamma range occurring during the presentation of the mind wandering and creativity task stimuli are specific to the DMN.

### DMN dynamics

The tasks employed in the current study encouraged semantic and mind wandering processes over a long time-window. The previous results focused solely on the initial DMN engagement (averaged over the first 2 seconds of the stimulus stage). To fully leverage the task design, we investigated the dynamics of theta and gamma in DMN over a long time-course (0-15 seconds, in 29 time-bins), focusing on both the stimulus encoding (Figure 3B) and the response stage of the tasks (Figure 3C). First, we verified that task type (AUT, MW), response stage (Stimulus, Response), and their interaction were important for both theta and gamma models (sequential model comparisons adding each fixed effect and interaction term: all p<0.001; all post-hoc comparisons p-adj<0.001; average theta power across the 15s ± standard error for AUT-Stimulus: 137.4±4.9%; MW-Stimulus:143.2±4.9%; AUT-Response: 125.9±4.6%; MW-Response: 102.8±4.3%; gamma for AUT-Stimulus: 24.5±1.3%; MW-Stimulus: 20.1±1.2%; AUT-Response: 30.8±1.8%; MW-Response: 35.5%2.5).

Next, we tested for the effect of time. The dynamics were different across tasks and stages, denoted by the importance of the effect of time-bin and its interaction with the combinations of the tasks and stages (Figure 3B-C; model comparison with/without time-bin in theta: χ^2^(1)=248.3, p<0.001; gamma: χ^2^(1)=49.7, p<0.001; model comparison with/without the interaction terms between time-bin and the other fixed effects of task type and response stage in theta: χ^2^(3)=31.6, p<0.001; gamma: χ^2^(3)=151.3, p<0.001). During the stimulus processing stage (lighter colored lines in Figure 3B), there is a larger increase in gamma power for AUT than for MW, sustained for the first 6-8 seconds. In theta, we see a similar effect but in the opposite direction, with a relative decrease in theta power modulation during the first 6-7 seconds of AUT; after ∼7 seconds, AUT-related theta modulation climbs back up to the same level as for MW. This temporal evolution potentially reflects a more critical initial stimulus processing stage during AUT, for which the semantic information about the object needs to be searched, ideas generated and compared to previous experiences in order to evaluate alternative uses for that object. During the response stage (darker colored lines in Figure 3C), DMN engagement is stronger for MW than for AUT, with a greater decrease in theta and greater increase in gamma over the course of the response window. The enhanced DMN recruitment during response production in the MW task could be reflecting the increased memory demands of recalling and reporting a train of thought.

### Dorsomedial and lateral DMN locations contribute differently to mind wandering and creativity

In the previous sections, we showed that DMN is recruited in both mind wandering and creativity, albeit in different ways. In addition, we found that distinct DMN subregions exhibit differential engagement during task execution. In a model comparing lateral temporal and dorsomedial DMN subsystems, the effect of DMN-subsystem identity and its interaction with the other variables (task type, task stage and time) was significant (model comparison with/without DMN-subsystems and their interaction with the other fixed effects in theta: χ^2^(8)=267.03, p<0.001; gamma: χ^2^(8)=392.7, p<0.001). Given the complexity of the full model (effect of task type, task stage, DMN-subsystem, time, and their interactions), we performed post-hoc comparisons using a simplified model (refitted without the interaction between time and the other variables), and uncovered the following 3-way interaction: theta power was differentially modulated across the two DMN subsystems during mind wandering, but not AUT: theta power was higher in *dorsomedial* DMN (vs. lateral DMN) during the MW stimulus stage, but higher in *lateral* DMN (vs. dorsomedial) during the MW response stage (Figure 4A; lateral vs. dorsomedial DMN during MW-Stimulus: z=-6.6, p-adj<0.001; lateral vs. dorsomedial DMN during MW-Response: z=2.6, p-adj=0.037; during AUT-Stimulus: z=-1.7, p-adj=0.37; during AUT-Response: z=-2.1, p-adj=0.14). Meanwhile, gamma power was higher in the lateral DMN (vs. dorsomedial) during the response stage of *both* tasks (lateral vs. dorsomedial DMN during MW-Stimulus: z=1.95, p-adj=0.2; lateral vs. dorsomedial DMN during MW-Response: z=8.1, p-adj<0.001; during AUT-Stimulus: z=2.4, p-adj=0.06; during AUT-Response: z=4.4, p-adj<0.001). The opposite theta modulation in dorsomedial and lateral DMN during the stimulus vs. response stages of MW indicates that the process of mind-wandering recruits different portions of the DMN depending on the ongoing cognitive process (internalized mind wandering versus externalized-response production stage). No such dissociation was present in the gamma band, as gamma power was consistently higher in the lateral DMN regions, especially during the response stage, for both tasks. These results are suggestive of different sub-specializations within the DMN. To comprehensively unravel the complex interplay between the neural signatures occurring within the DMN for each task type and stage observed so far, we performed a hierarchical clustering analysis based on the similarity of the neural dynamics, described below.

**Figure 4.**
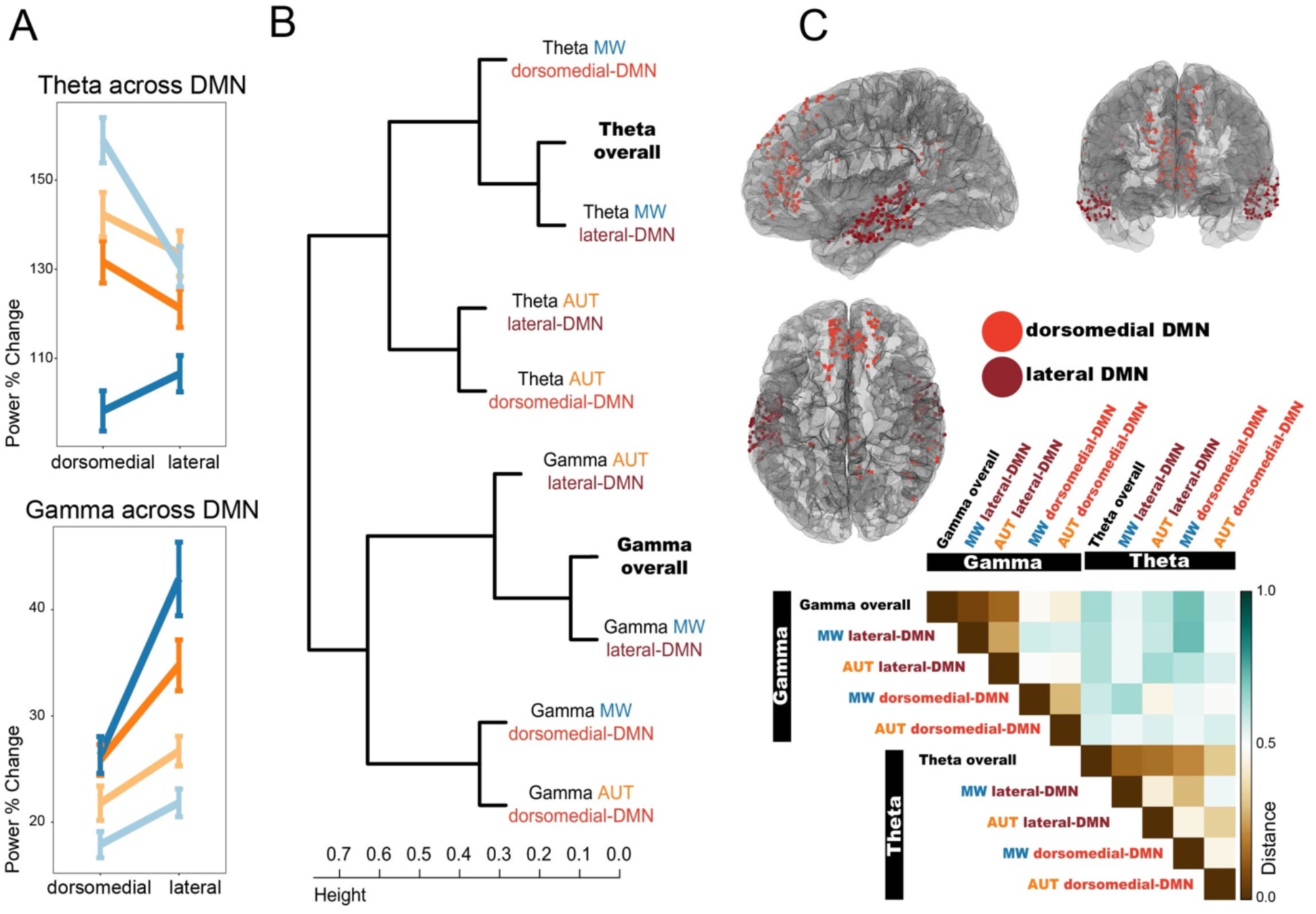
Interplay between theta and gamma dynamics and spatial contributions within the DMN. Panel A: spatial distribution of the task and stage interaction effects within the DMN. Theta power differences across tasks and stages are most prominent in dorsomedial electrode locations within the DMN, while gamma differences are larger in lateral temporal locations. Panel B: Hierarchical clustering results obtained by evaluating the distance between the vectors of observations formed by the time-course of power values for each stage (stimulus, response) and subject (limiting the analysis to n=10 subjects with electrodes in both DMN subsystems. Panel C top: location of electrodes recording from the DMN, color coded by the anatomical subdivision in two subsystems. Dorsomedial-DMN includes medial and dorso-lateral frontal-parietal areas (electrodes recording from ventromedial prefrontal cortex, anterior cingulate, superior frontal gyrus, posterior cingulate, postero-medial parietal, parahippocampal gyrus). Lateral-DMN includes lateral temporal areas (electrodes recording from middle temporal gyrus, superior and middle temporal sulci). Panel C bottom: Average distance matrix across variables. Panels B and C bottom highlight the different direction of theta and gamma modulations during the tasks (first cluster separation in B, highest distance values in C) and illustrate how the spatio-temporal features of theta and gamma provide non-redundant information about the ongoing cognitive processes. Theta can be used to separate the tasks, while gamma separates the DMN subsystems (second clustering layer: theta separates MW from AUT; gamma separates lateral-DMN from dorsomedial-DMN). In addition, the second clustering layer provides insight on the similarity of the overall gamma and theta effects (averaged across tasks and DMN nodes), suggesting that overall theta is mostly resembling theta during MW, while overall gamma modulations are mostly driven by lateral-DMN.

### Interplay between neural signatures and DMN nodes

The degree of similarity between the temporal evolution of theta and gamma activity during each task for the two DMN subsystems was assessed using hierarchical clustering. In this analysis, we also included overall theta and gamma dynamics (regardless of the subsystem or specific task) to help identify the main drivers of the overall neural patterns during creativity and mind wandering (Figure 4B). The first main separation occurs between theta and gamma, demonstrating that the activity in these two frequency bands display opposite patterns of modulation, where relative decreases in theta power are paired with increases in gamma activity (large distance values denoting anticorrelation in Figure 4C bottom; electrodes for each DMN subsystem in 4C top). In addition, the two frequency bands display different similarity patterns within their subclusters. Theta power can be used to separate the two tasks (0.96 bootstrap probability, b.p.). Indeed, theta power is similar across the two subsystems, and mainly resembles the whole DMN activity during mind wandering (0.86 b.p.). In contrast, gamma activity can be used to differentiate dorsomedial from lateral DMN subsystems (0.95 b.p.). Gamma is different across the two DMN subsystems, being overall driven by the lateral temporal DMN nodes (1.00 b.p.), while displaying similar dynamics across tasks. Overall, this result demonstrates that theta and gamma play differential, non-redundant roles during mind wandering and creativity, and display different spatial distribution within the DMN.

### High-frequency stimulation of electrodes within the DMN reduces creativity

Finally, we assessed whether the creativity of the responses during mind-wandering and alternate uses was causally dependent on DMN activity. To this end, we manipulated DMN activity by delivering high-frequency stimulation over a pair of electrodes located in the DMN. We evaluated the behavioral data from the subjects that received stimulation (n=9) and compared the originality scores for stimulation versus non-stimulation trials (Figure 5). Stimulation had the effect of reducing originality in AUT (Figure 5A-B; median SemDis without stimulation: 0.97; with stimulation: 0.94; W=75, p=0.001), while it had no effect over mind wandering (Figure 5C; SemDis without/with stimulation: 0.97 vs. 0.97; W=28, p=0.3). We performed a control analysis on the AUT effect by including the subjects that did not receive stimulation (n=4, median SemDis: 0.97). The control analysis confermed that the reduction in the AUT-originality scores can be reliably detected in a larger sample (i.e., by including trials from patients that did not receive stimulation and testing for differences between stimulated and non-stimulated trials in the full sample, n=13; W=2638.5, p=0.017).

**Figure 5.**
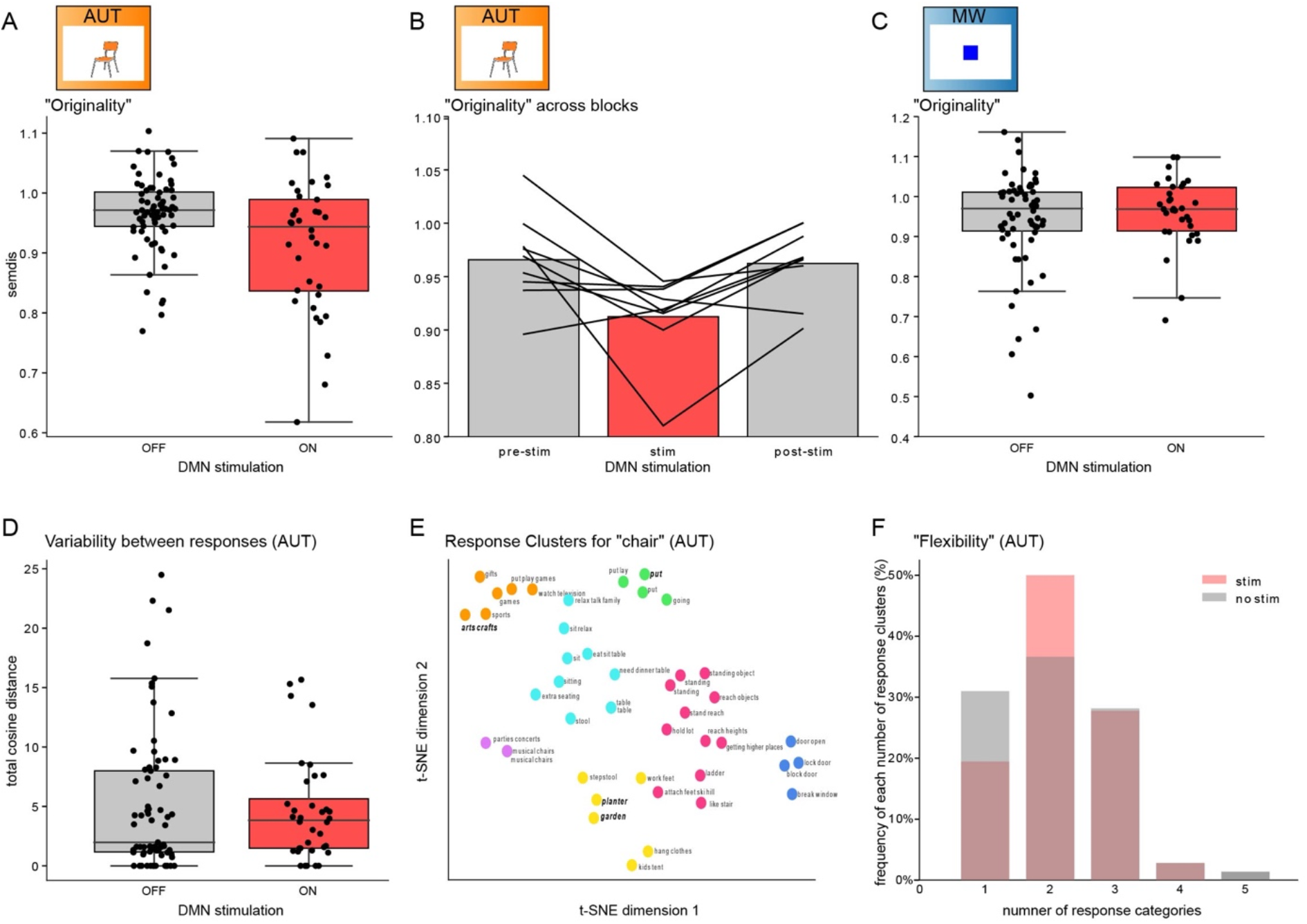
Stimulation of DMN reduces creativity of responses. Panel A: SemDis scores (y axis) during AUT as a function of stimulation. The SemDis scores, a proxy of the originality of the responses, are reduced during stimulation, indicating a decrease in the originality (each point represents an AUT item score across 9 subjects). Panel B: Same data as panel A showing the subject-level effect of stimulation (each line represents a participant). Panel C: SemDis scores during MW are not modulated by stimulation. Panel D: Total cosine distance, a proxy of variability between the responses, was not affected by stimulation. Panel E: Example responses to the item “chair” across the first two t-SNE dimensions, color coded by the semantic category clusters. Responses from one participant are highlighted in bold to showcase the number of semantic clusters spanned by that participant. Panel F: Histogram indicating the number of categories into which each participant’s responses for each item can be categorized. The number of response clusters, an index of flexibility, was not affected by stimulation.

Our observation that stimulation during AUT diminished SemDis scores suggests a decrease in the originality of the responses: the responses during stimulation were more semantically similar to the cue word itself. But does DMN stimulation also modulate other aspects of creativity, such as fluency and flexibility? A naïve test of fluency simply counts the number of responses to each presented item. However, multiple similar responses may be “original” (semantically different from the cue word), but virtually identical to each other, so may inflate the “fluency” count for that trial. To address this confound, we devised an alternate metric of fluency that also includes the *variability* of the responses—consequently, this altered metric of fluency can be called “variability”. Specifically, this metric considers the number of responses weighted by how distinct they are from each other, such that very similar responses would not contribute as strongly to this altered metric of fluency. Notably, unlike the originality score above, this analysis does not account for the identity of the cue (item) itself, nor the distance between it and the responses. We quantified the relationship between each pair of responses as the cosine distance between the two response vectors; the sum of the distances for all response pairs (total cosine distance) was used as the measure of variability. In our paradigm, stimulation did not affect the variability (Figure 5D; mean total cosine distance without stimulation: 5.04; with stimulation: 4.59; Wilcoxon signed-rank test, p = 0.65. Another facet of creativity is *flexibility*, which quantifies the number of distinct categories into which the responses fall. In order to analyze the effect of stimulation on flexibility, we performed t-SNE clustering on the entire set of responses for each item, and then used an affinity propagation clustering algorithm to identify semantic categories. Figure 5E shows the outcome of this clustering for one example item (“chair”). Then, for each patient, we can count the number of clusters from which that patient’s responses came. The number of response categories per item ranged from 1-5 (Figure 5F). For example, the responses of the patient highlighted (bold/italic lettering) in Figure 5E come from 3 clusters (even though the patient gave 4 responses total). Comparing the number of response categories across patients for trials with and without stimulation yielded no significant results (Wilcoxon signed rank test, p = 0.91), suggesting no effect of DMN stimulation on flexibility (Figure 5F).

## Discussion

In this paper, we utilized network-wide intracranial sampling of the DMN in order to dissect fast temporal dynamics underlying its function. We used LFP-derived spectral power changes in theta and gamma bands to differentiate network activation during a DMN-only state (mind-wandering task, MW) from creativity (alternate uses task, AUT). Our goal was to investigate how these very different modes of thought engage the DMN. First, we demonstrated network activation during both of our DMN-recruiting tasks, evidenced by anti-correlated activity between high and low-frequency power bands. This activation was DMN-task specific and did not occur during our visual attention control task, or outside of the DMN in a control network. Next, we examined the time course of DMN activation in the alternate uses vs. mind wandering tasks. We observed an early increase in gamma power in the alternate uses task, suggesting that the initial presentation of the object for which the subject needs to find possible alternate uses “triggers” DMN activity; in contrast, the mind wandering task led to later DMN recruitment, indicating that the network is more active during the retrieval and verbalization of the train of thoughts. Following this general investigation of DMN activity as a whole, we looked into DMN sub-networks and showed that the dorsomedial and lateral temporal areas are differentially engaged by our experimental tasks. Lastly, we probed the causal relationship between DMN activity and creative thinking, and we demonstrated that DMN inhibition through direct cortical stimulation reduces the originality of the responses without impairing other domains of creativity. In addition, mind wandering was unaffected by stimulation, suggesting that it is either less dependent on DMN integrity, or that AUT is more dependent on the DMN ability to synchronize with other networks.

### Theta and gamma-band signals are modulated in opposite directions

Neural activity within the DMN was modulated during periods of trial presentation versus pre-trial baseline periods. If we compare the DMN tasks to the control attention task, we find an increase in gamma (30-70Hz) and *relative* decrease in theta power (4-8Hz) specifically for the DMN tasks (AUT and MW). This kind of anti-correlation between low and high frequency is a classic feature of large scale neural recruitment and network engagement across various systems, related to the balance between excitation and inhibition, and often characterized as a shallower slope of the power spectrum (Gao et al 2017).We did not observe this pattern in a non-DMN control network (somatomotor network). From these findings, we can conclude that alternate uses and mind wandering tasks successfully engaged DMN, compared to a control task (ATT) and a non-DMN network (somatomotor).

### Task-dependent temporal evolution of DMN recruitment

When considering the initial DMN activity during the stimuli presentation stage, AUT displayed the strongest effect (compared to MW and to the control ATT), with the greatest relative decrease in theta power and the greatest increase in gamma activity (Figure 3A). This result was based on the first 2 seconds of AUT and MW in order to offer a time-matched comparison to our control attentional task (ATT); looking across the *entire* time course revealed a dichotomy between the temporal dynamics of DMN activity in the AUT and MW tasks. Specifically, the temporal evolution of DMN activity indicates that the alternate uses task engages the network in the *early* stage of the task (Figure 3B), while participants are presented with an item and need to covertly search through semantic and episodic information for alternative uses. During the later response selection stage, the theta range pattern is quite similar, while gamma activity is even stronger (Figure 3C). In contrast, mind wandering displayed a weak response in the early task stage and a very strong effect during the later response production stage, when the participant is asked to recollect out loud their train of thoughts (Figure 3C). Given previous findings (Kajimura et al 2016, Vallat et al 2022) of DMN involvement in mind wandering, the relatively lower DMN recruitment during the MW stimulus period is surprising: the stimulus period of the MW task is the phase during which the actual process of “mind wandering” occurs – during the response window, the thoughts are merely reported, and this response more likely represents episodic memory related DMN activation. However, during the stimulus period, mind wandering was not entirely spontaneous, as participants were also actively viewing a colorful shape – perhaps the stimulus acted as a “distraction” from the mind wandering aspect, drawing the participants’ attention and making it harder for them to detach from the visual stimulus and allow for spontaneous thought to take over. Indeed, there is evidence that external sensory input can inhibit mind wandering (Christoff 2012). Moreover, the experimental setting in which the patients are sitting in a hospital room, postoperatively, may also impair their ability to mind wander. It is also noteworthy that most of the evidence on mind wandering arises from fMRI studies (Fox et al 2015), thus slow fluctuations (<4Hz, often <0.1Hz), not examined here but known to modulate theta and gamma in DMN nodes (Foster & Parvizi 2012), may offer a more direct correlate of the mind wandering process (Kucyi et al 2018). Altogether, this evidence suggests that different cognitive functions elicit the DMN with their unique signatures of network activation, warranting further investigation to dissect the specificity of these function-signals relations.

Another significant implication of our result stems from the relative importance of the stimulus in the AUT vs MW tasks: for the creativity task, the stimulus plays a critical role, guiding the semantic and episodic search for alternate uses. During the mind wandering task, on the other hand, the stimulus is just a probe to relax, thus initiating the spontaneous mind wandering process, with no explicit instruction, nor requirement to perform any memory-guided processes. When the response window begins, however, participants engage in memory retrieval to report their train of thought. From this perspective, the DMN activity reported in our findings seems to track the memory-intensive portions of the task, in line with current views that semantic and episodic memory systems are highly integrated with DMN activity (Menon 2023). Our findings isolate the key role played by active recollection of memories as a driver of DMN activity, either as a covert search of alternative uses, or as overt retrieval of a previous train of thoughts.

### Spatial distribution of theta and gamma modulations within the DMN

So far, we discussed the dynamics of DMN activity as a whole, but the effects we observed differed across regions of the DMN. In particular, theta effects displayed a complex 3-way interaction between tasks, their stages and DMN subnetworks: lateral temporal cortex was the site exhibiting the major modulation (i.e., relative decrease in power) for creativity. Mind wandering on the other hand presented a dissociation between stimulus stage effects – stronger in lateral-DMN, like AUT – and response stage, featuring an especially strong decrease in theta power in dorsomedial DMN. Meanwhile, gamma modulation was clearly driven by lateral temporal DMN, with a similar pattern for mind wandering and creativity during both task stages (Figure 4A).

To assess the spatio-temporal features of theta and gamma more holistically, we decided to exploit the temporal evolution of the signals without the artificial separation in two stages, revealing their associations using similarity-based analysis. We demonstrate that theta power differentiates mind wandering from creativity, displaying stronger modulation during mind wandering. Thus, theta power dynamics featured strong dissimilarities between our tasks. By connecting the similarity analysis results with the 3-way interaction results discussed above (Figure 4A-B), theta may be orchestrating changes in the cognitive process and requirements, signaling the recruitment of lateral temporal cortex during early-stage mind wandering and throughout creativity. The engagement of dorsomedial areas (PCC, SFG, mPFC) during the recollection stage of the mind wandering process could be reflecting memory retrieval, as theta-phase synchronization between posterior medial cortex and non-DMN regions has been previously implicated in autobiographical judgements (Foster et al 2013).

Gamma-band signals, classically considered to be reflecting local neural activity, are distinct between the DMN subsystems, but quite similar across the tasks: lateral temporal structures are the major contributors to the observed gamma power effects for both AUT and MW tasks (Figure 4). Our findings demonstrate that theta and gamma signatures provide non-redundant information about the ongoing DMN activity: theta is modulated differently according to the cognitive processes at play, while gamma is recruited in a similar manner by all processes, but with a strong spatial gradient biased toward lateral temporal cortex. The steady contribution of lateral temporal gamma activity across our tasks is congruent with this region’s implication in inner speech and semantic processing. Indeed, the proximity of DM and language areas within the lateral temporal cortex is highly suggestive of multiple adjacent distributed networks that work closely to support different facets of the same higher cognitive function (DiNicola & Buckner 2021), like the ability to construct and maintain an internal narrative, continuously ongoing during mind wandering. Semantic processing and category-specific memory, which are key components of the AUT task, have been historically localized to the middle temporal gyrus, a subregion of the lateral temporal DMN (Moore & Price 1999). Furthermore, the MTG has been associated with the ability to attribute an action (use) to an object (Martin & Chao 2001), a cognitive process certainly involved in the alternate uses task. In fact, MTG seems particularly attuned to categories related to “tools” (Perani et al 1999), matching our AUT stimuli. Beyond internal speech and semantic processing, the lateral temporal cortex is also important for memory. Closed-loop stimulation of the lateral temporal cortex has been employed to enhance episodic memory, (Ezzyat et al 2018), implying that memory performance does not rely uniquely on the classic medial temporal network, but that later-temporal DMN subnetworks make distinct and behaviorally-relevant contributions. By considering our results in this context, we corroborate the modern view of the DMN as an heterogenous and multiplexed system, comprised of anatomo-functional subnetworks subserving different cognitive functions (Menon et al 2022).

### Causal relationship between DMN activity and creativity

Creativity is a multifaceted concept that can have multiple meanings in different contexts. For divergent thinking, as tested by the alternate uses task, the creativity of responses can be quantified in several ways. For instance, novel, surprising responses that are very distinct from the traditional use of the cue object would score high in *originality*. On the other hand, *fluency* refers to the ability to produce a large number of responses. However, since fluency does not traditionally consider the identity of the responses, if a participant gives many nearly-identical responses, the fluency score will be high. Thus, we designed an alternative measure to better reflect a participant’s ability to “fluently” produce a variety of responses. This metric, called *variability*, is similar to fluency in that it is higher for a larger number of responses; but it also takes into account how distant each response is from all the other responses. As a result, similar responses are less strongly weighted than distinct responses. Finally, another variation of fluency is *flexibility*—or the number of semantic categories of the responses. Like variability, flexibility also down-weights similar responses and amplifies distinct responses. However, it does so in an “all-or-nothing” manner: all responses that fall within the same semantic category are treated as a single response.

Having defined these aspects of creativity, we asked whether the DMN is involved in any of them; if so, does the DMN play an equal role in the different facets of creativity, or do originality and fluency differentially depend on DMN activity?

Previous work using direct brain stimulation suggests that dominant, lateral DMN nodes contribute to creative fluency, but not originality (Shofty et al.,2022). Yet, in the current study, disrupting DMN activity with high-frequency stimulation reduced only the originality of the responses, but had no effect on fluency, variability, or flexibility of the content. In other words, stimulation specifically modulated how original the alternative uses produced were, without affecting the variability across responses or number of semantic categories used (Figure 5). There were notable differences in the behavioral scoring choices and stimulation locations that could explain these different findings. Shofty et al. only included responses that were substantially different from the common use of each object, whereas the current study counted all responses. For the originality measure, responses other than the traditional use of the object were also excluded if they were frequently produced, to avoid inflating the fluency. In the current study, DMN stimulation occurred in different locations across subjects; either medial or dorsal PFC and, for one subject only, in lateral temporal cortex. In contrast, Shofty et al. stimulated the lateral dominant DMN exclusively. The partial effect on creativity reported here (as well as the partial, though different, effect in Shofty et al.) may therefore be related to the different anatomical target. Based on our electrophysiology results, the lateral-temporal DMN seems to be a more critical hub, as this subsystem was the main driver of local neural activity (gamma) for AUT (as well as MW). This interpretation aligns well with existing literature about the effect of stimulation on memory and free recall: for instance, Ezzyat et al. (2018) demonstrated that stimulation in the lateral temporal region of the DMN was able to robustly modify memory performance (Ezzyat et al 2018). It is possible that the effect on originality reported in our study reflects the reduced ability to access memories related to the objects, consequently affecting the ability to generate original and non-canonical responses, without interfering with other aspects. More research will be needed to disentangle the causal contribution of different DMN subsystems to the widespread set of cognitive processes that underlie divergent thinking.

Besides the alternate uses task, we also examined the effect of DMN stimulation on mind wandering. The trains-of-thought reported in the mind wandering task were unaffected by stimulation, suggesting that creativity processes rely more strongly on DMN integrity than mind wandering. Our results indicate that the thought processes occurring during day-dreaming and mind wandering are more resilient to external perturbation of the system. It is possible that changes in the train of thoughts might be more difficult to detect given their “unconstrained” fluctuating nature, also in line with the subjective experience of a constant and coherent internal narrative. In addition, we scored the mind wandering responses using a similar approach to the one used for alternate-uses, while it is possible that the cognitive process that occurred during our MW task could not be appropriately measured through semantic distance metrics. Another possible explanation for the lack of perturbation effects on mind wandering originates from our electrophysiology findings: MW displayed a complex interaction between task stages and theta activity across the two DMN subsystems. Theta is typically regarded as a long-range communication channel; therefore, to affect mind wandering, it might be necessary to tap into lower frequency communication channels with patterned stimulation (while the current study used high frequency stimulation) and potentially stimulate several DMN nodes simultaneously.

### Limitations and future directions

Although standard for human invasive experiments, our study is based on a small sample size (n=13). To limit potential bias, we employed within-subject approaches for all our analysis, and all group-level analysis are based on the consistency of within-subject differences. By nature of the technique employed we were also limited to clinically-defined electrode trajectories, constraining our ability to equally sample within and across individual cortical areas. For example, the PCC, a key DMN hub was rarely sampled in our cohort. Consequently, our subdivision in dorsomedial and lateral DMN is an oversimplification of the network and its subsystems, arising in part from previous evidence and in part by our cortical sampling availability. In particular, our dorsomedial-DMN system spanned medial and dorsal frontal areas, posterior medial cortex and parietal locations, potentially mixing together in one subsystem several unique contributions. This said, our study was not aimed at dissecting the individual contribution of each DMN node, and whole-brain techniques are better suited to inform on the precise organization of largely distributed subnetworks.

There are very few studies employing direct cortical stimulation of DMN regions during active behavior (Foster & Parvizi 2017, Shofty et al 2022). In general, stimulation experiments are particularly challenging and require the consideration of several clinical factors, often constraining the ability to perform rigorous controls (i.e., selecting consistently the same stimulation site across individuals, repeating the experiment with a control stimulation site outside the network or sham condition). In this study, we were unable to select posterior medial cortical locations as the stimulation sites, preventing us from drawing direct comparisons with previous studies targeting PCC. Our results demonstrate that the originality of the response given during a creativity task can be diminished by interfering with different nodes of the DMN. More studies will be needed to understand the *necessary* intra- and inter-network communication mechanisms and how these relate to findings of juxtaposed of neural populations with episodic and executive functional profiles (Aponik-Gremillion et al 2022). Indeed, creativity and divergent thinking has also been shown to involve cognitive control networks such as the executive control network (Beaty et al 2014, Benedek et al 2014, Ellamil et al 2012, Gonen-Yaacovi et al 2013). For example, studies have demonstrated the significance of key brain regions associated with top-down control, such as frontoparietal system, in the performance of a variety of cognitive tasks linked to creativity (Abraham et al 2012, Fink et al 2009). Creativity seems to be subsisted by multiple brain networks, such as the executive and the salience network, and by their synchronization with the DMN to support the generation of novel, divergent thoughts (Beaty et al 2015, Beaty et al 2016, Beaty et al 2018).

## Acknowledgments

We thank all the patients that took part in the experiments. We thank Brian Metzger for his assistance in the early stages of the project and Sandy Reddy for data pre-processing support. This work was supported by the McNair Foundation (S.A.S.) and NIH (R01-MH127006 to K.R.B.).

## Author contributions

K.R.B. S.A.S. B.S. designed research; E.D. H.Q.D. R.K.M. B.R.P. A.A. J.A. performed research; E.B. E.D. H.Q.D. R.R. analyzed data; E.B. E.D. R.R. K.R.B. S.A.S. B.S. wrote paper.

